# Differential Antibody Recognition by Novel SARS-CoV-2 and SARS-CoV Spike Protein Receptor Binding Domains: Mechanistic Insights and Implications for the Design of Diagnostics and Therapeutics

**DOI:** 10.1101/2020.03.13.990267

**Authors:** Ilda D’Annessa, Filippo Marchetti, Giorgio Colombo

## Abstract

The appearance of the novel betacoronavirus SARS-CoV-2 represents a major threat to human health, and its diffusion around the world is causing dramatic consequences. The knowledge of the 3D structures of SARS-CoV-2 proteins can facilitate the development of therapeutic and diagnostic molecules. Specifically, comparative analyses of the structures of SARS-CoV-2 proteins and homologous proteins from previously characterized viruses, such as SARS-CoV, can reveal the common and/or distinctive traits that underlie the mechanisms of recognition of cell receptors and of molecules of the immune system.

Herein, we apply our recently developed energy-based methods for the prediction of antibody-binding epitopes and protein-protein interaction regions to the Receptor Binding Domain (RBD) of the Spike proteins from SARS-CoV-2 and SARS-CoV. Our analysis focusses only on the study of the structure of RBDs in isolation, without making use of any previous knowledge of binding properties. Importantly, our results highlight structural and sequence differences among the regions that are predicted to be immunoreactive and bind/elicit antibodies. These results provide a rational basis to the observation that several SARS-CoV RDB-specific monoclonal antibodies fail to appreciably bind the SARS-CoV-2 counterpart. Furthermore, we correctly identify the region of SARS-CoV-2 RBD that is engaged by the cell receptor ACE2 during viral entry into host cells.

The data, sequences and structures we present here can be useful for the development of novel therapeutic and diagnostic interventions.

## Introduction

The outbreak of the new betacoronavirus 2019-nCoV, formally named SARS-CoV-2, represents a world-wide epidemic threat that has been causing major disruptions in societies and economies^1-2^. For these reasons the new virus has been internationally declared a public health emergency.

In this context, structural knowledge of the proteins of the virus can provide important information to develop strategies against the infection, ranging from diagnosis to therapy. In general, coronaviruses use this protein localized on the envelope to bind their cell receptors. After binding, a series of molecular events lead to the fusion of the viral membrane with that of the host cell, eventually leading to virus entry and invasion.

One of the necessary steps in this mechanism is the binding of the spike protein Receptor Binding Domain (RBD) to the cell receptor Angiotensin-Converting Enzyme 2 (ACE2). Indeed, knock out of ACE2 in HeLa cells was shown to prevent infection.

For these reasons the development of antibodies that target the spike RBD of coronaviruses and block their interaction with ACE2, disrupting a fundamental infection mechanism, represent attractive therapeutic solutions. Monoclonal antibodies with high affinities had been previously developed against the spike RBD of SARS-CoV. However, despite the correlations between the two viruses, no RBD-targeted SARS-CoV monoclonal antibodies have shown the ability to bind and neutralize 2019-nCoV.

The recent publication of the Cryo-EM structure of the 2019-nCoV spike trimer at 3.5 Å resolution can provide a useful basis to link these observations to molecular causes, guide the design of SARS-CoV-2 specific antibodies, and possibly suggest the basis for molecules able to block cell viral entry.

Here, we carry out a comparative analysis of the spike protein RBDs from SARS-CoV-2 and SARS-CoV to reveal the key properties of their surfaces that can be reconnected to observed differences in antibody (Ab) and receptor binding. Starting from the atomic resolution information available from the solved structures of the two proteins, the MLCE (Matrix of Low Coupling Energies) method is applied to identify the subsets of surface residues that can be defined as "interacting". MLCE analysis of protein energetics accounts for the interactions that each residue establishes with all other residues of the protein it belongs to. The correlation of this parameter with the structural properties of the protein thus highlights which substructures are (pre)organized to interact with a potential partner (a cell receptor or an antibody).

MLCE analyzes the interaction energies of all of the amino acids in a protein. In particular, it computes the nonbonded part of the potential (van der Waals, electrostatic interactions, solvent effects) via an MM/GBSA calculation, obtaining, for a protein composed by *N* residues, an *N* × *N* symmetric interaction matrix *M_ij_*. Eigenvalue decomposition of the matrix highlights the regions of strongest and weakest couplings: the fragments that are on the surface, contiguous in space and weakly coupled to the protein core, define the potential interaction regions. In other words, the putative interacting patches are assumed to be characterized by frustrated intramolecular interactions. The actual interaction with a partner will actually occur if favorable interactions determine a lower free energy for the bound than the unbound state^3-4^. Indeed, Minimal energetic coupling with the rest of the protein allows these substructures to undergo conformational changes, to be recognized by a binding partner (Antibodies, Receptors). All these properties are hallmarks of Antibody binding epitopes and PPI substructures.

The MLCE method for the prediction of Protein-Protein Interaction regions and Antibody binding epitopes has been validated in several previous studies^3-14^.

## Results and Discussion

Here, we took advantage of the recent publication of the structures of SARS-CoV-2 spike protein in its pre-fusion state (doi.org/10.1126/science.abb2507) and of known structure of the SARS-CoV spike protein to comparatively apply our interaction epitope prediction method MLCE.

The application of MLCE to the Receptor Binding Domain (RBD, aa 319-591) of the SARS-CoV-2 spike protein (pdb code <u>6vsb</u>; doi: <u>10.2210/pdb6vsb/pdb</u>)^15^ and of the SARS-CoV spike protein (pdb code <u>5×58</u>; doi: 10.2210/<u>pdb5X58</u>/pdb) **(Figure 1)** shows that the two protein regions that contain potential interaction regions/Ab-binding epitopes share the same topological localization at the two opposite ends of the 3D structures **(Figure 1)**. However, this trait appears to be the only common one. The linear sequences that correspond to the predicted interacting epitope regions are reported in **Table 1.**

**Figure 1.**
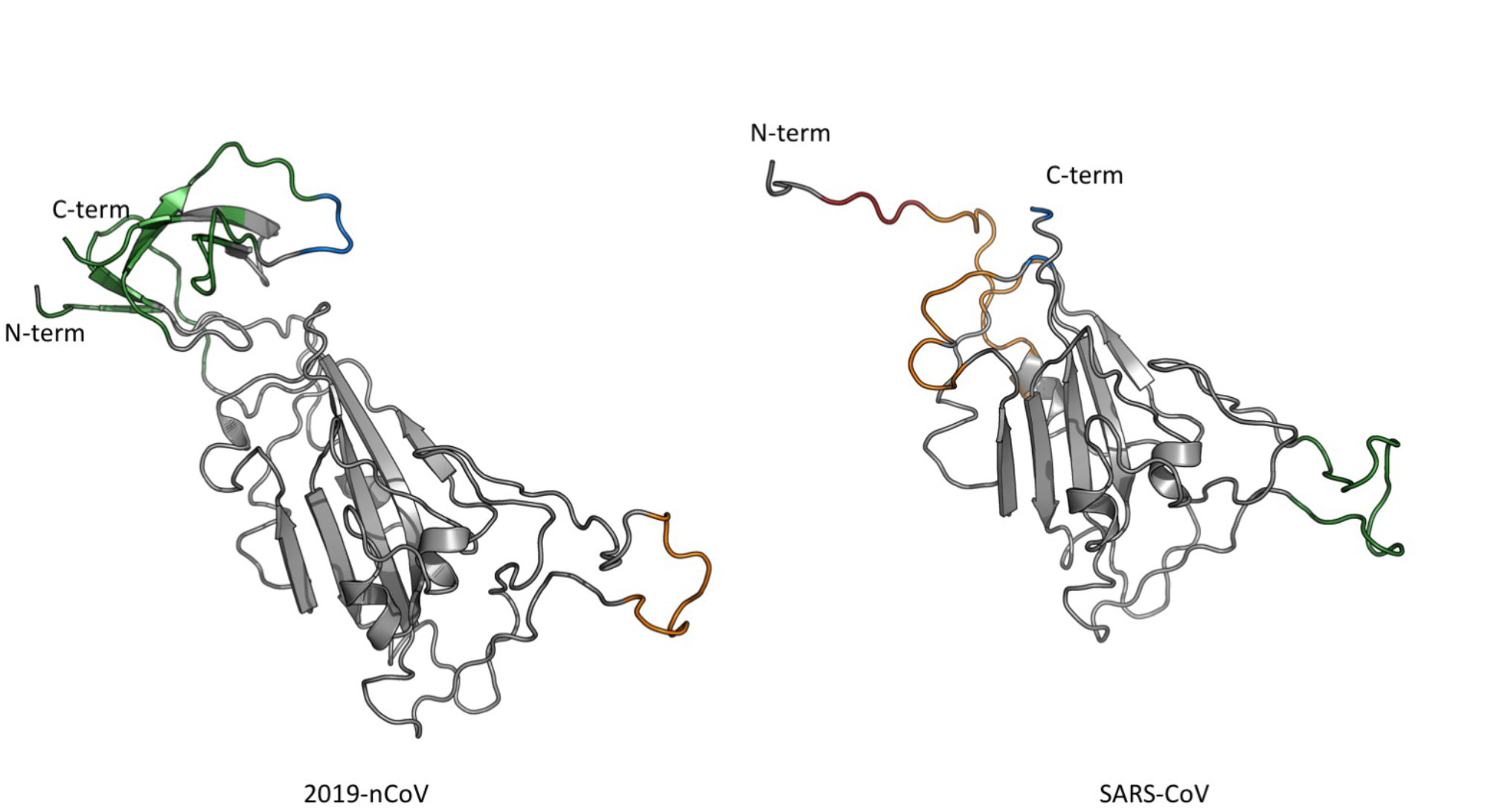
Comparison of the Receptor Binding Domains of the SARS-CoV-2 and SARS-CoV spike proteins. The predicted epitopes, whose sequences are reported in Tables 1 and 2 are highlighted with colors.

**Table 1.**
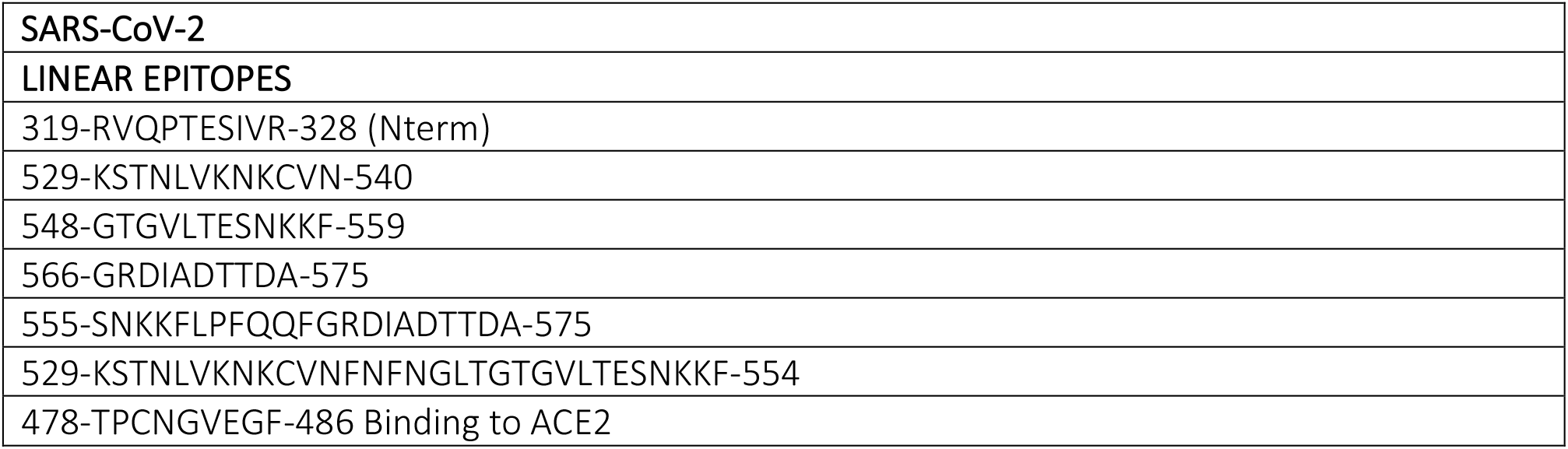
Epitope sequences for the Receptor Binding Domain (RBD, aa 319-591) of the SARS-CoV-2 spike protein (pdb code <u>6vsb</u>).

|  |
| --- |
| **SARS-CoV-2** |
| **LINEAR EPITOPES** |
| 319-RVQPTESIVR-328 (Nterm) |
| 529-KSTNLVKNKCVN-540 |
| 548-GTGVLTESNKKF-559 |
| 566-GRDIADTTDA-575 |
| 555-SNKKFLPFQQFGRDIADTTDA-575 |
| 529-KSTNLVKNKCVNFNFNGLTGTGVLTESNKKF-554 |
| 478-TPCNGVEGF-486 Binding to ACE2 |

Importantly, the predicted epitopes for SARS-CoV spike RBD show a good degree of overlap with the epitopes mapped for neutralizing Ab 80R^16^, S230^17^ and m396^18^, providing a relevant validation of our approach. However, in spite of the high degree of structural similarity between the two spike proteins, these mAbs failed to bind SARS-CoV-2 RBD, probably due to the low degree of sequence conservation in the regions contacted by these antibodies. The fact that most of the SARS-CoV-directed mAbs are not reactive against the novel betacoronavirus once more highlights the need to discover specific epitopes in SARS-CoV-2.

Actually, except for the common topological localization, the predicted Ab-binding epitopes differ between the two proteins, both in their sequence and 3D structural organization **(Table 1, 2 and Figure 1)**. In some cases, linear sequences spanning distant segments of the primary structure come together in the three-dimensional structure to form extended substructures potentially (pre)organized for binding. This is particularly true for the conformational epitope at the N- and C-terminal region of SARS-CoV-2 RBD: the interacting region spans a large part of a well-defined betasheet subdomain **(Table 1, 2 and Figure 2)**. It is tempting to suggest that the expression of such subdomain in isolation could give rise to a highly immunoreactive molecule^19-21^.

**Figure 2.**
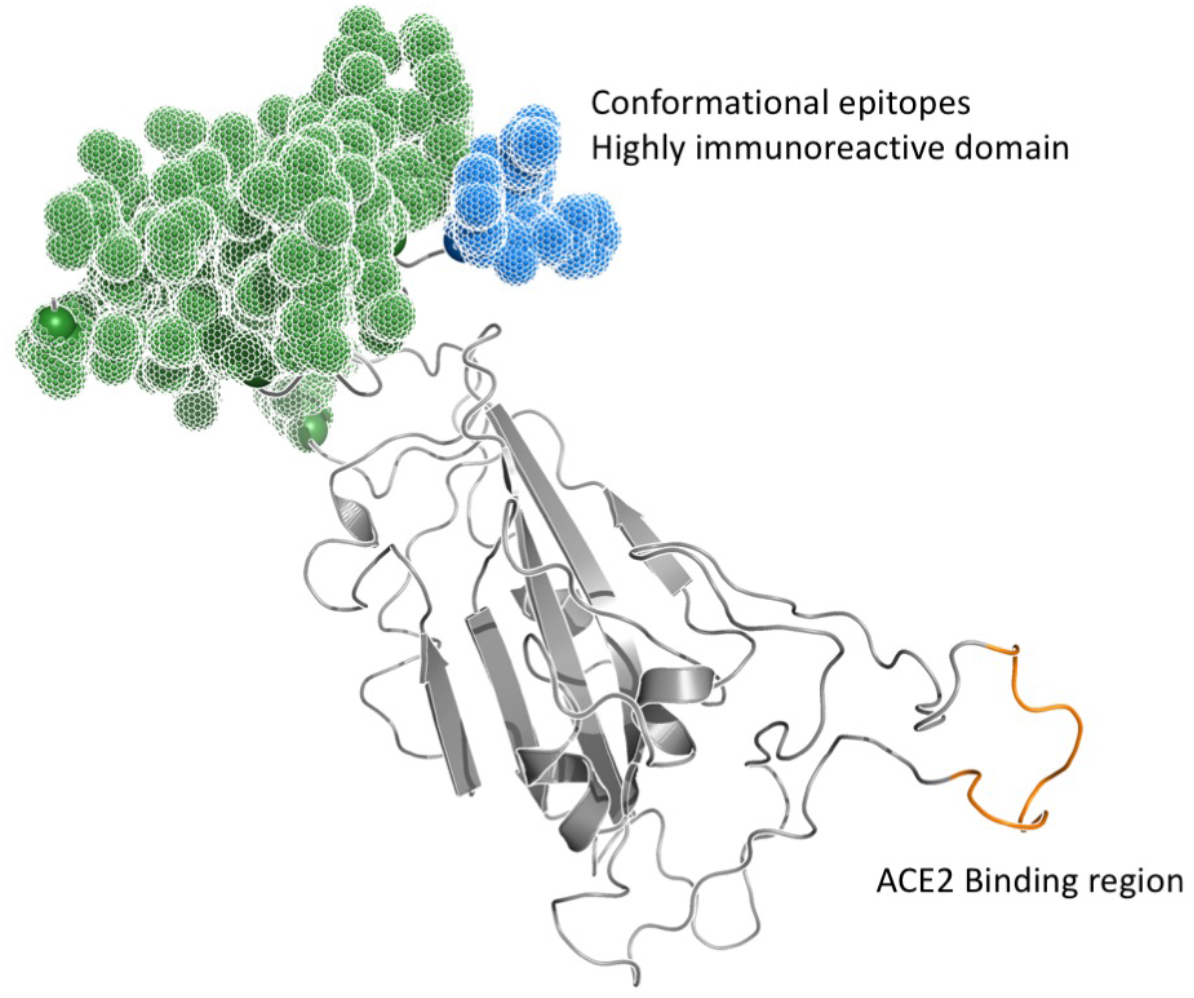
The space filling regions indicate possible conformational epitopes for SARS-CoV-2 formed by the juxtapositions of the sequences reports in Table 1. The region binding to ACE2 and correctly predicted is also highlighted.

**Table 2.** Epitope sequences for the Receptor Binding Domain (RBD, aa 319-591) of the SARS-CoV spike protein (pdb code <u>5×58</u>).

|  |
| --- |
| **SARS-CoV** |
| **LINEAR EPITOPES** |
| 315-RFPNITNLCPFG-326 |
| 370-SATKLNDLCFS-380 |
| 458-VPFSPDGKPCTPPALNCYWPLN-479 Binding to ACE2 |

In SARS-CoV-2 RBD, the knowledge of substructures representing potential conformational epitopes may be helpful in guiding the development of peptide-based immunogens able to elicit antibodies that specifically target the cognate protein to which the predicted epitopes belong (**Figure 2**). Simply speaking, the termini of the linear segments making up the conformational epitopes could be aptly bridged with a number of Gly residues sufficient to approximate the distance in the experimental structure (Table 3). Antibodies raised against these molecules, which recapitulate the main determinants of immunoreactivity of the full protein, could have useful applications in a therapeutic setting^19-21^. Furthermore, information on potentially specific SARS-CoV-2 spike RBD-specific epitopes can guide the synthesis of mimics of the epitope sequences in the form of peptides for use in diagnostic settings. Serological approaches aimed at revealing antibodies elicited in the host in response to the infection or/and at using specifically selected antibodies to detect relevant 2019-nCoV antigens could indeed be used for the screening of people who may have come into contact with the virus as well as for epidemiological surveillance, a relevant issue considering the number of asymptomatic patients which caused a fast and widespread diffusion of SARS-CoV-2. In general, serological approaches of this kind can be simple and quick to perform and could aptly represent a valid complement to swabs followed by molecular nucleic acid-based methods, currently used as the front-line response to detect the virus in the window period in which host B-cell responses have not yet been elicited.

**Table 3.** Proposed sequences for the mimics of the conformational epitopes predicted for the Receptor Binding Domain (RBD, aa 319-591) of the SARS-CoV-2 spike protein (pdb code 6vsb). Boldface G’s indicate glycines inserted to bridge two parts of a conformational epitope; Bold-face, underlined S’s or T’s indicate serines inserted to substitute for original cysteines or methionines, respectively.

|  |
| --- |
| **CONFORMATIONAL EPITOPES for SARS-CoV-2** |
| 320-VQPTES -325-**GGG**-529-KSTNLVKNK**S**VN-541-**GG**-548-GTGVLTE-553 |
| 555-SNKKFLPFQQFGRDIADTTDA-575-**GG**-I-**GG**-587-ITP**S**S-591 |
| 566-GRDIADTTDA-575-**GG**-I-**GG**-587-ITP**S**S-591 (Cterm) |

The specific SARS-CoV-2 RBD epitope sequences and structures identified here can thus be used to design new molecular probes (immunoprobes) with facile synthetic accessibility and the potential to function both as antigen-mimicking baits to target virus-specific antibodies in patients’ fluids, and as immunogens to generate new Abs able to detect relevant antigens^22-23^.

Importantly, after we ran our predictions on the SARS-CoV-2 RBD, crystal structures of this domain in complex with the cellular receptor protein ACE2 was reported by Lan et al. (aa 480-488)^24^ and by Yan et al.^25^. Of note, one predicted interaction region almost completely overlaps with the substructure of spike protein RDB engaged in binding with ACE2 (478 TPCNGVEGF 486). Synthetic mimicry of this sequence could aptly generate high priority candidates aimed at disrupting PPIs between SARS-CoV-2 RBD and its cell receptor ACE2. Such molecules could be used to block viral entry into cells.

Similarly, our approach also predicts a patch on SARS-CoV spike RBD, which is partly involved in binding ACE2 (see^26^).

Overall, while the two RBD proteins likely evolved from a common ancestor and share a common fold, they are characterized by different sequences. The physico-chemical properties determined by such sequence differences, efficaciously unveiled by MLCE, reverberate in the modulation of the networks of intraprotein interactions that underlie the (pre)organization of regions for potential recognition of binding partners (cell receptors or antibodies).

These results represent, to the best of our knowledge, one of the first molecular physical-chemistry based rationalization of the reasons why some of the tested SARS-CoV RBD monoclonal Abs do not have appreciable cross-reactivity to SARS-CoV-2 RBD^15^, despite the fact that the two domains were recently reported to display the same affinity for the ACE2, with K_D_ values of 15.2 nM and 15 nM for SARS-CoV-2 and SARS-CoV, respectively^27^.

In this complex scenario, our predictions can help disentangle some of the issues related to betacoronavirues infectivity and immunological response.

## Acknowledgement

We acknowledge funding by Regione Lombardia, project READY (Regional Network for developing diagnostic methods in rapid response to emerging epidemics and bio-emergencies)

## Methods

The coordinates of the proteins analyzed through the text were downloaded from the pdb with the following codes:

2019-nCoV spike glycoprotein Receptor Binding Domain: PDB ID <u>6vsb</u>

SARS-nCoV spike glycoprotein Receptor Binding Domain: PDB ID <u>5×58</u>

Missing residues belonging to highly flexible loops not solved in the structure of the Spike protein were modeled by homology using the SwissModel web server (https://swissmodel.expasy.org/), that retrieved the structure of the SARS-CoV <u>5×58</u> as the best template.

The structures were used as input for epitope prediction using the Matrix of Local Coupling Energies method (MLCE), which combines the analysis of structural determinants of a given protein with its energetic properties^3-4^. This approach allows to identify nonoptimized, low-intensity energetic interaction-networks, corresponding to those substructures that can be more prone to establish interactions with Antibodies, and be suitably recognized by binding partners. Briefly, the contiguous regions on the protein surface that are deemed to have minimal coupling energies with the rest of the structure are selected on the basis of the eigenvalue decomposition of the matrix reporting the non-bonded interaction of all residue-pairs. The eigenvector associated to the most negative eigenvalue permits to reconstruct a simplified matrix which reports the maximal and minimal stabilizing residue-pairs in the protein structure. Filtering of the simplified matrix with the contact matrix allows to identify contiguous residue-pairs characterized by their essential degree of coupling to the rest of the protein. Selection of proximal pairs showing minimal coupling with the rest of the protein defines putative epitopes. Selection is carried out on the basis of a threshold value (called softness), which defines the percentage of the set of putative interaction sites by including increasing residue-residue coupling values until the number of couplings that correspond to the lowest contact-filtered pairs under the threshold was reached.

## Detailed Prediction of protein interaction surfaces: MLCE

MLCE is a technique based on the analysis of the interaction energies of all the amino acids in a protein^3, 28-31^. In particular, it computes the non-bonded part of the potential (van der Waals, electrostatic interactions, solvent effects) via a MM/GBSA calculation, obtaining, for a protein composed by *N* residues, a *N × N* symmetric interaction matrix *M_ij_*. This matrix can be expressed in terms of its eigenvalues and eigenvectors as

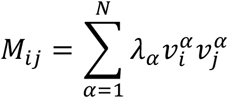

where *λ_a_* is the *α*-th eigenvalue and 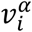 is the i-th component of the corresponding eigenvector. The eigenvector with the most negative correspondent eigenvalue contains most of the interaction information for the stabilizing interaction of the system. An approximated interaction matrix 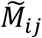 is thus given by

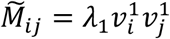

If the structure of the protein is known, one can estimate a contact matrix *C_ij_* by assuming two amino acids in contact if the distance between two of their heavy atoms is smaller than a threshold. The Hadamard product of the two matrices gives us the matrix of the local coupling energies

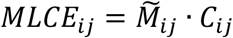

We will select as possible interacting zones sets of close by residues that show weak or frustrated interactions.

The analysis of the energetic properties of the surface residues is based on the MLCE method. Basically, we perform a MM/GBSA analysis of the structure in a force field, obtaining a symmetric per-residue interaction matrix *M_ij_* keeping only non-bonded interaction (i.e. electrostatic, van der Waals and solvation contributions). We diagonalize the matrix, obtaining a set of eigenvectors *x*^(*i*)^ sorted following the increasing value of their eigenvectors *λ_i_* where *N* is the number of amino acids in the sequence. We thus can write the original matrix *M_ij_* as

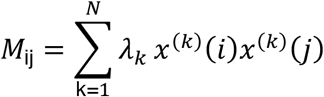

It has been shown that the first eigenvector alone can be used to build an approximate interaction matrix *M*: *M*_ij_=*λ*_1_*x*^(1)^(*i*)*x*^(1)^(*j*), which recapitulates the interactions most relevant for the stabilization of a certain conformation of a defined protein or protein substructure.

The MM/GBSA is made with Amber 14 software with the ff14SB forcefield.

## Notes

#### Summary of Updates

The title was changed (upon suggestion by a reader) to better highlight the applicative implications of our mechanistic studies. We also added a table of designs to better explain the gly-bridging concept we suggested for conformational epitopes.

